# CD28-Selective Inhibition Prolongs Non-Human Primate Kidney Transplant Survival

**DOI:** 10.1101/2023.05.03.539333

**Authors:** Brendan P. Lovasik, Steven C. Kim, Laura Higginbotham, Walter Wakwe, David V. Mathews, Cynthia Breeden, Alton B. Farris, Christian P. Larsen, Mandy L Ford, Steven Nadler, Andrew B. Adams

## Abstract

Costimulation blockade using belatacept results in improved renal function after kidney transplant as well as decreased likelihood of death/graft loss and reduced cardiovascular risk; however, higher rates and grades of acute rejection have prevented its widespread clinical adoption. Treatment with belatacept blocks both positive (CD28) and negative (CTLA-4) T cell signaling. CD28-selective therapies may offer improved potency by blocking CD28-mediated costimulation while leaving CTLA-4 mediated coinhibitory signals intact. Here we test a novel domain antibody directed at CD28 (anti-CD28 dAb (BMS-931699)) in a non-human primate kidney transplant model. Sixteen macaques underwent native nephrectomy and received life-sustaining renal allotransplantation from an MHC-mismatched donor. Animals were treated with belatacept alone, anti-CD28 dAb alone, or anti-CD28 dAb plus clinically relevant maintenance (MMF, Steroids) and induction therapy with either anti-IL-2R or T cell depletion. Treatment with anti-CD28 dAb extended survival compared to belatacept monotherapy (MST 187 vs. 29 days, p=0.07). The combination of anti-CD28 dAb and conventional immunosuppression further prolonged survival to MST ∼270 days. Animals maintained protective immunity with no significant infectious issues. These data demonstrate CD28-directed therapy is a safe and effective next-generation costimulatory blockade strategy with a demonstrated survival benefit and presumed advantage over belatacept by maintaining intact CTLA-4 coinhibitory signaling.

## Introduction

Renal transplantation is the treatment of choice for end-stage renal disease (ESRD) demonstrating improvements in patient quality of life, patient survival, and cost effectiveness compared to hemodialysis(1–3). Despite the significant advances in post-transplant short-term survival, several studies have highlighted that long-term outcomes remain less than optima(4). Standard immunosuppression maintenance agents such as calcineurin inhibitors (CNI) in part contribute to late renal allograft failure through direct effects including acute and chronic nephrotoxicity, interstitial fibrosis and tubular atrophy, focal glomerular sclerosis as well as indirect effects including higher rates of hypertension, diabetes mellitus, and increased cardiovascular risk (5, 6). In addition, the inability to control the alloimmune response resulting in the development of anti-donor antibody either due to immunosuppressive drug minimization or discontinuation also plays an important role in late allograft failure. In an effort to improve post-transplant outcomes there was a concerted effort to develop immunosuppressants that more specifically target T cells with the goal of achieving fewer off-target effects compared to calcineurin inhibitors (7). Belatacept, a costimulatory blockade agent, was approved by the Food and Drug Administration (FDA) in 2011. Long-term results of the BENEFIT study, a Phase III clinical trial of belatacept in kidney transplant recipients, demonstrated that its use led to improved patient/graft survival, significant increases in renal graft function compared to cyclosporine (8) as well as improvements in blood pressure and serum lipid profiles (9). Unfortunately higher rates and grades of acute rejection within the first year post-transplant have prevented its widespread clinical adoption (9).

Treatment with belatacept, which binds CD80 and CD86 on antigen-presenting cells (APC), blocks both positive (via CD28) and negative (via CTLA-4) T cell signaling. CD28-selective therapies may offer improved potency by blocking CD28-mediated costimulation while allowing CTLA-4 coinhibitory signals to reach the T cell (10). Allowing ligation of CTLA-4 decreases T cell activation and proliferation (11, 12). Furthermore, CTLA-4 expression contributes to the proper suppressive function of regulatory T cells (Tregs) with reports of fatal autoimmune disease and decreased Treg function *in vitro* and *in vivo* in CTLA-4 knockout mice (13). Prior efforts to translate targeted CD28 therapy to human clinical trials were halted when the superagonist mAb TGN1412 resulted in cytokine storm and multi-organ failure in six healthy human volunteers in a 2006 Phase I clinical trial (14). Accordingly, any new anti-CD28-based therapy will require evaluation of agonistic activity. In support of CD28-directed therapy, Liu et al. demonstrated that use of a selective anti-CD28 domain antibody (dAb) prolonged skin graft survival to >100 days compared to treatment with CTLA4-Ig (median survival time [MST] 34 days) in a mouse model (15). Reports of immunosuppression with an antagonist anti-CD28 monovalent Fab’ antibody fragment, FR104, in nonhuman primate (NHP) renal transplantation and arthritis models also exist (16, 17). Suchard et al. recently reported the development of a novel domain antibody directed at the human CD28 receptor, anti-CD28 dAb or BMS-931699 (lulizumab) (18). They evaluated the anti-CD28 dAb for agonistic activity and found no change in expression of activation markers, proliferation, or cytokine production after exposure of T cells to anti-CD28 dAb *in vitro* (18). Anti-CD28 dAb was also evaluated in 180 healthy subjects and there were no significant cytokine or immune cell changes noted over a variety of doses used (13). More recently, an additional 275 patients with systemic lupus erythematosus were treated with anti-CD28 dAb in a Phase II clinical trial with a favorable safety profile (19). Given the previous preclinical data and recent clinical trial evaluation of CD28-directed therapies it seemed reasonable to test the effectiveness of anti-CD28 dAb in our nonhuman primate (NHP) model of kidney transplantation. Initial study efforts focused on *in vitro* comparisons of anti-CD28 dAb to belatacept. Successful *in vitro* results led to examination of anti-CD28 dAb as a monotherapy regimen in a NHP model of renal transplant. To model a clinically translatable immunosuppressive protocol incorporating anti-CD28 dAb, two additional experimental groups were included that incorporated standard maintenance immunosuppressive agents (mycophenolate mofetil (MMF) and steroid) with either anti-IL-2R mAb or T cell depletion for induction. Anti-CD28 dAb was significantly more potent than belatacept in preventing allograft rejection when compared directly as a monotherapy. The inclusion of anti-CD28 dAb in a clinically relevant immunosuppression regimen led to long-term survival of kidney allografts supporting its evaluation as a next-generation costimulatory blockade therapy in transplant patients.

## Materials and Methods

### T Cell Stimulation Assay

Peripheral blood mononuclear cells (PBMCs) were collected from naive Rhesus macaques and isolated from CPT Tubes (BD Biosciences) after centrifugation. PBMCs (2.5×10^5^ cells per well) were incubated in 96-well flat-bottom plate and stimulated at a ratio of 2:1 in the presence of NHP-specific MACSiBead™ Particles loaded with biotinylated antibodies to mimic antigen-presenting cells (T Cell Activation/Expansion Kit, Miltenyi Biotec). Responder cells were labeled with cell trace violet prior to incubation. Cells proliferated for seven days at 37°C and 5% CO_2_ in R10 medium with IL-2 (100 ng/mL). During incubation, responder PBMCs were stimulated in the presence of belatacept or, anti-CD28 dAb as indicated. Cell proliferation was assessed by cell trace violet dilution on flow cytometry.

### Renal transplantation

Twenty adolescent Rhesus macaques underwent bilateral native nephrectomy followed by life-sustaining renal allotransplantation using a kidney from a fully MHC-mismatched donor (Yerkes National Primate Research Center, Emory University, Atlanta, GA, USA). Operations were performed in accordance with Institutional Animal Care and Use Committee regulations. Postoperatively, graft function was assessed by daily monitoring of urine production and weekly serum laboratory evaluations. Peripheral blood was collected for weekly cellular subset analysis using flow cytometry. Quantitative cytomegalovirus (CMV) titers were measured weekly using a polymerase chain reaction (PCR)-based assay. Clinically significant CMV viremia was defined as >10,000 copies per mL, and oral antiviral therapy was initiated when positive. Ultrasound-guided renal biopsies were performed postoperatively at days 35, 70, and 140. Biopsy specimens were submitted for hematoxylin and eosin (H&E) staining (Yerkes National Primate Research Center, Emory University, Atlanta, GA, USA). A blinded renal pathologist performed histologic analysis. Endpoint criteria for euthanasia were defined as (A) serum creatinine >5 mg/dL or blood urea nitrogen (BUN) >100 mg/dL on two consecutive measurements or (B) rejection-free survival to 600 days.

### Treatment regimen

Animals were assigned to one of four treatment arms: anti-CD28 dAb monotherapy (n=6), belatacept monotherapy (n=5), or anti-CD28 dAb, MMF, steroid taper and anti-IL-2R mAb induction (n=5), or anti-CD28 dAb, MMF, steroid taper and T cell depletion induction (n=4). Group A recipients received anti-CD28 dAb (3 mg/kg) monotherapy beginning 1 day prior to transplantation with weekly injections until post-operative day (POD) 70, biweekly injections until POD 140, and then monthly injections until POD 364 (*Figure 1*). Anti-CD28 dAb is attached to polyethylene glycol (PEG) for prolonged half-life and administered as a subcutaneous (sc) injection on a weekly or biweekly basis (17). Group B recipients received Belatacept (20 mg/kg) infusions administered intravenously (IV) on POD 0 and POD 4, biweekly until POD 56, and then monthly until POD 140. Group C received anti-CD28 dAb and in addition anti-IL-2R mAb (basiliximab) induction (POD 0 and 4) with daily MMF/steroids through POD 140 (*Figure 1*) and was notably free of CNI. Group D received anti-CD28 dAb, MMF, and steroid as described for group C and received T cell depleting therapy for induction instead of anti-IL-2R mAb. Anti-CD28 dAb and belatacept were obtained from Bristol-Myers Squibb (New York, NY, USA). T-cell depletion was accomplished using anti-CD4 and anti-CD8 mAbs. T-cell depletion began 1–3 days prior to transplantation with peri-transplant dose(s) of anti-CD4 50 mg/kg IV (clone CD4R1), and anti-CD8 50 mg/kg IV (clone M-T807R1; both from NIH Nonhuman Primate Reagent Resource). Basiliximab, MMF, and steroids were purchased from McKesson Corporation (San Francisco, CA, USA).

**Figure 1.**
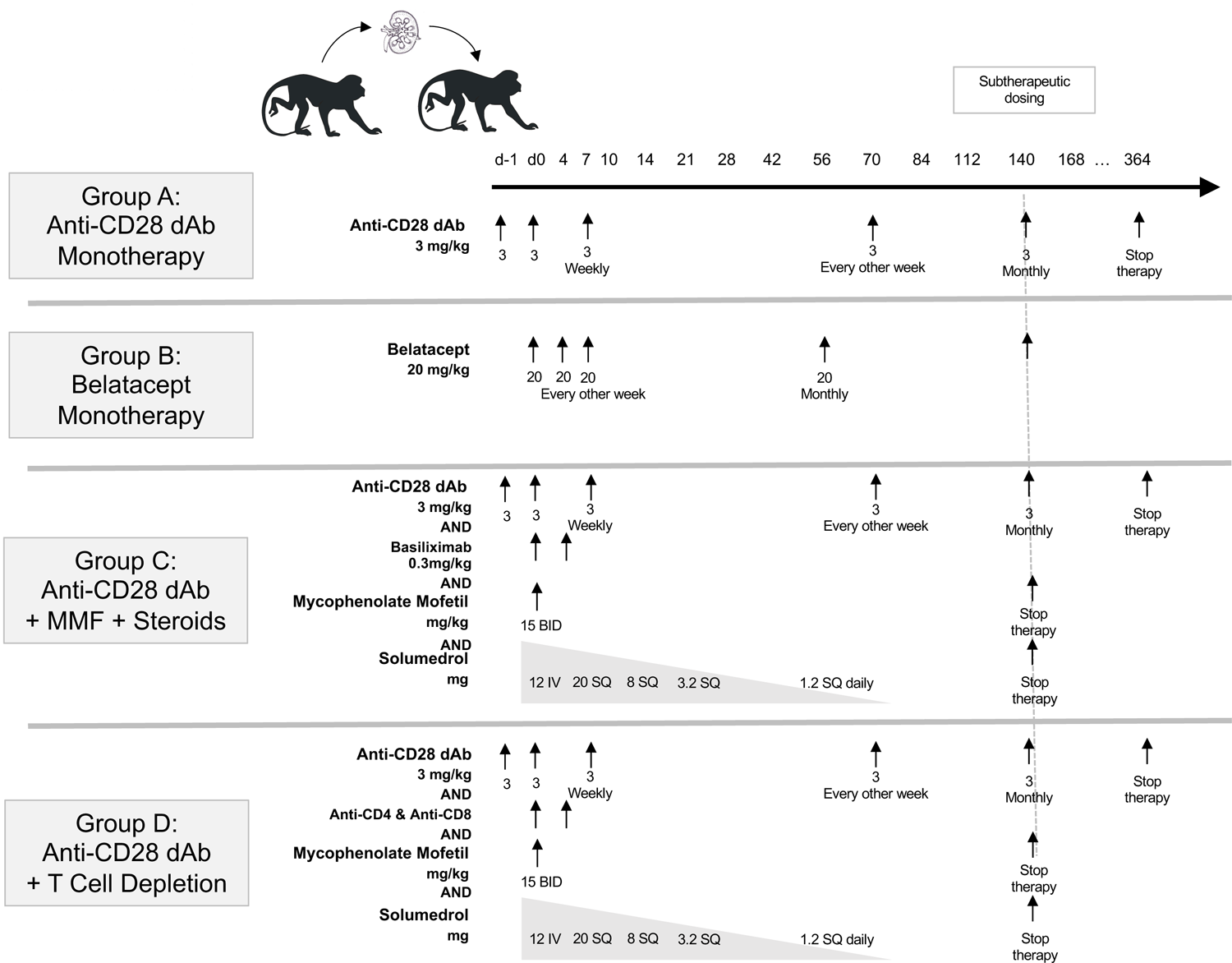
CD28 dAb Dosing regimen. (A) Anti-CD28 dAb was received as a subcutaneous injection weekly POD 0-70, biweekly POD 70-140, and monthly POD 168-364. Therapy was discontinued after POD 364. (B) For the anti-CD28 dAb plus conventional immunosuppression treatment arm, anti-CD28 dAb was administered as above. Recipients in the adjunct conventional immunosuppression treatment arm received basiliximab induction at POD 0 and 4, while recipients in the T lymphocyte depletion received preoperative doses of anti-CD4 and anti-CD8 depleting antibodies as described in the Methods section. Animals in both arms also received daily MMF and steroids until POD 140.

### Pharmacokinetics of anti-CD28 dAb in Rhesus macaques

Peripheral blood samples were collected from the femoral vein of recipients pre-treatment then weekly post-transplant for pharmacokinetic analysis. Drug level measurements were performed by Bristol-Myers Squibb.

### Flow cytometry

Peripheral blood was collected pre-transplant and weekly post-transplant to assess the response of immunologic cellular subsets to CD28-directed therapy. Cell surface staining was performed using fluorescent antibodies against human CD3 (clone SP34-2), CD4 (552838), CD8 (RPA-T8), CD45RA (556626), and CCR7 (150503) from BD Biosciences (San Jose, CA, USA); and anti-human CD28 (CD28-2) and CD25 (BC96) from eBioscience (San Diego, CA, USA).

Intracellular fixation and permeabilization was performed using the eBioscience (San Diego, CA, USA) buffer set in accordance with manufacturer’s instructions. Intracellular staining was completed using antibodies against human Foxp3 (PCH101, eBioscience), Helios (22F6, Biolegend, San Diego, CA, USA), and CTLA-4 (557301, BD Biosciences). Samples were collected on a BD Biosciences LSR II flow cytometer and analyzed using FlowJo software, version 10.0.7 (Treestar Inc., Ashland, OR, USA).

### Donor-specific antibodies

Peripheral blood samples were obtained from donor Rhesus macaques at time of donor nephrectomy to collect and store PBMC, as previously described. Blood from renal transplant recipients was collected weekly and spun to collect serum. Recipient serum from POD 0, 14, 28, 56, 84, 112, 150, 200, 300, 400, and 500 was incubated with thawed donor PBMC to determine presence of donor-specific antibodies (DSA). Staining was performed using goat anti-monkey IgG and IgM conjugated to Alexa-488 (KPL; Gaithersburg, MD, USA). Results were collected on a BD Biosciences LSR II flow cytometer with analysis of antibody binding by FlowJo version 10.0.7 (Treestar Inc). Results were reported using MFI (mean fluorescence intensity).

### Statistical analyses

Post-transplant survival times were plotted and compared using Kaplan-Meier survival curves and log-rank tests with p-values <0.05 considered significant. All analyses were performed using GraphPad Prism, version 6.0 (La Jolla, CA, USA).

## Results

### Anti-CD28 dAb inhibits NHP T cell proliferation in vitro

Selective blockade of the CD28 pathway inhibits T cell activation by preventing CD28 engagement with CD80 and CD86 while preserving the ability of CTLA-4 to bind to these identical ligands. Previous reports have shown that targeting the CD28-CD80/86 pathway with selective anti-CD28 antagonists is highly effective at preventing T cell activation *in vitro* and *in vivo*. Here, we investigated a novel anti-CD28 domain antibody composed of a Fc-devoid CD28-specific single-chain Fv domain that specifically binds to and blocks CD28 signaling, leaving CTLA-4 co-inhibition intact.(18) First, we investigated in vitro the consequences of selective CD28 binding on T lymphocyte stimulation. PBMCs from rhesus macaque recipients were stimulated with allogeneic stimulators for 7 days in a T cell stimulation culture. After stimulation we observed significant T cell proliferation in the absence of immunosuppression as measured by CTV dilution in both CD4^+^ and CD8^+^ T cells (*Figure 2A*). The presence of belatacept in the culture medium resulted in minimal decreases in CD4^+^ T cell proliferation and almost no reduction in CD8^+^ T cell expansion. Alternatively, the presence of anti-CD28 dAb completely controlled CD4^+^ T cell proliferation to level of the unstimulated control and significantly impacted CD8^+^ proliferation as well, amounting to a superior immunosuppressive effect compared to belatacept (*Figure 2B*). The effects of the anti-CD28 dAb on T cell proliferation demonstrated a dose-dependent effect, with reduced proliferation at higher drug concentrations. Given these promising *in vitro* results, anti-CD28 dAb was translated *in vivo* into a preclinical non-human primate model of renal transplantation.

**Figure 2.**
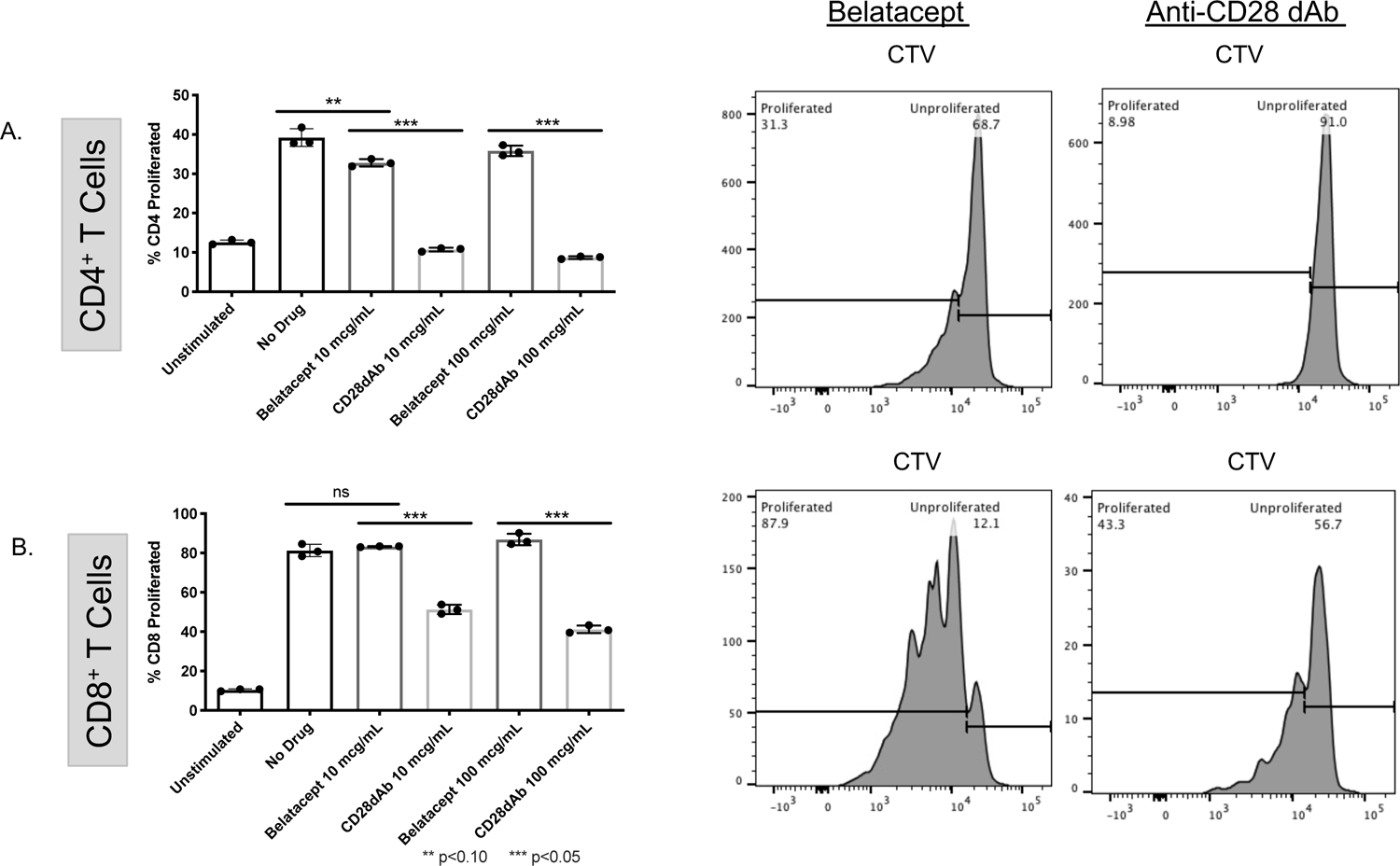
*In-vitro* inhibition of T cell proliferation with anti-CD28 dAb. Mixed Lymphocyte culture with NHP responder and stimulator cells treated with belatacept and anti-CD28 dAb at low (10mcg/mL) and high concentrations (100mcg/mL) (A) Stimulation of CD3^+^ T cells without immunosuppressive drug present in culture medium resulted in a 72.9% increase in relative proliferation from unstimulated controls. Proliferation is unaffected by treatment with belatacept at both concentrations. In contrast, treatment with anti-CD28 dAb resulted in a relative reduction CD3^+^ T cell proliferation of 68.2% (lo) and 76.5% (high) when compared to stimulated controls (p< 0.05 for both). Similar results for stimulation of CD4^+^ (B) and CD8^+^ (C) T cells where proliferation is unaffected by belatacept (B) CD4^+^ T cells stimulated in the presence anti-CD28 dAb exhibited reduced relative proliferation capacities of 72.5% (lo) and 79.5% (high) (p< 0.05 for both). (C) CD8^+^ T cells also exhibited reduced relative proliferation capacities of 36.6% (lo) and 50.1% (high) (p< 0.05 for both).

### Anti-CD28 dAb prolongs renal allograft survival in non-human primates and is synergistic with conventional immunosuppression

Dosing regimens for both anti-CD28 dAb and belatacept were scheduled to be intensive in the initial post-operative period when the risk of rejection is greatest, with the gradual extension of the interval between treatments over time eventually reaching subtherapeutic drug levels by day 140 (*Figure 1, Figure 3B*). In the anti-CD28 dAb treatment groups the early post-transplant phase consisted of weekly or biweekly injections, while the belatacept-treated recipients received biweekly infusions during the same time frame. Treatment with belatacept monotherapy prolonged survival compared to untreated animals who reject within the first week post-transplant (historic data) (*Table 1*). All belatacept-treated recipients rejected during the initial, high-intensity treatment phase. Monkeys treated with anti-CD28 dAb monotherapy demonstrated superior graft survival than belatacept monotherapy recipients, with median survival of 187 days compared to 29 days (*Table 1*). Within the anti-CD28 dAb monotherapy group, half of recipients experienced early rejection in a similar fashion to belatacept. However, the remaining three animals continued for 11-21 months on anti-CD28 dAb alone. One of these recipients rejected at 321 days after receiving subtherapeutic monthly injections for nearly 6 months (*Table 1, Figure 3A*). In contrast with the eventual and uniform rejection observed in belatacept monotherapy animals, two recipients treated with anti-CD28 dAb experienced indefinite survival after therapy was completely withdrawn at 1 year. Long-term graft survival was associated with relatively normal renal physiology as measured by serum creatinine *(Figure 4).* These animals were sacrificed at 600 days per clinical study protocol and exhibited no clinical or histologic evidence of rejection (*Supplemental Figure 1*).

**Figure 3.**
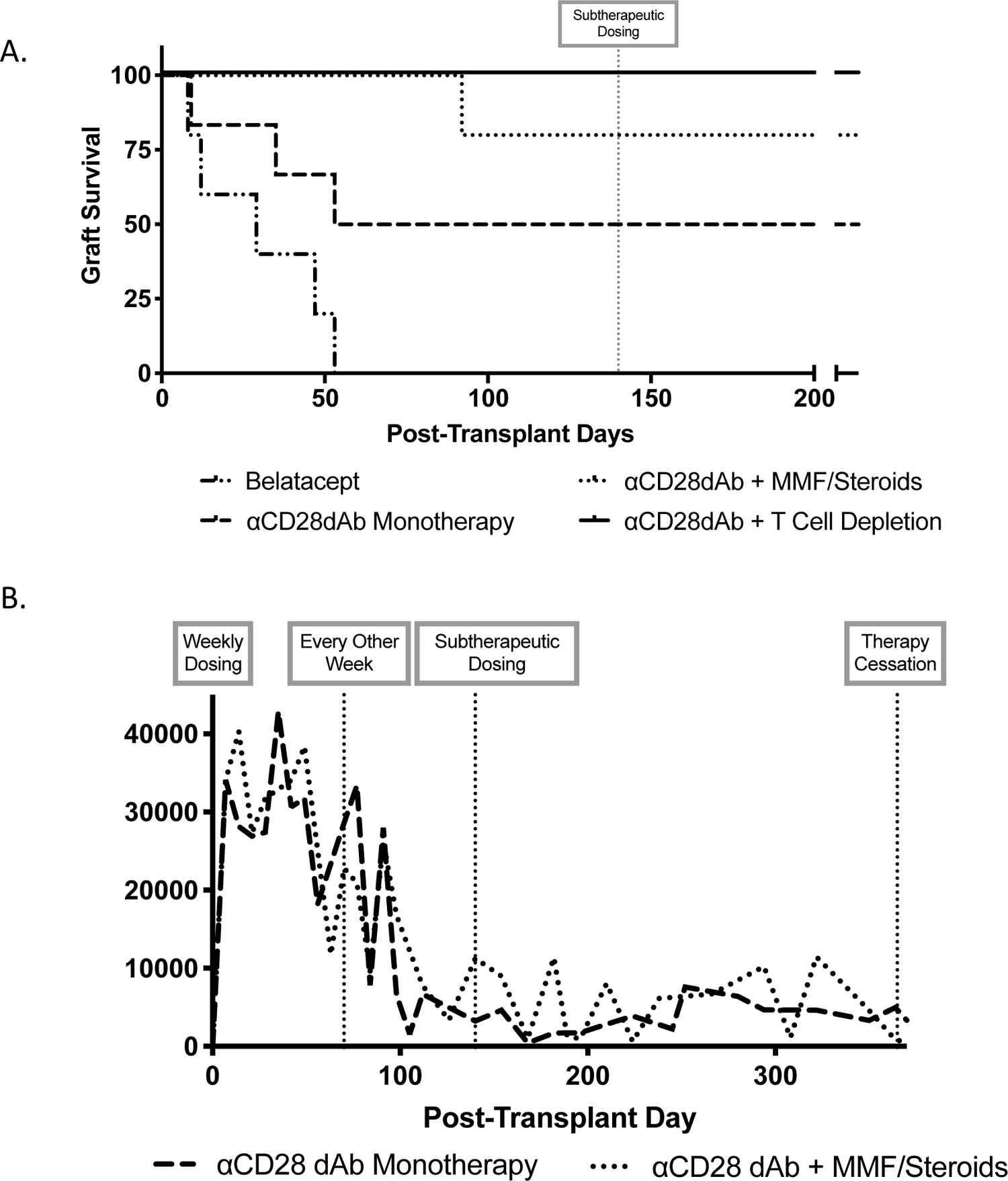
Survival after NHP renal transplantation. Treatment with anti-CD28 dAb prolongs survival in non-human primate renal transplantation. (A) All four regimens are compared. Vertical line indicates cessation of immunosuppressive therapy at POD 364. Six Rhesus macaques treated with anti-CD28 dAb alone survived >187 days MST compared to 29 days MST with belatacept alone, p=0.07. Two monkeys in the dAb-treated group remained ongoing at 600 days with rejection-free survival. Maximum survival in the belatacept-treated group was 53 days. Anti-CD28 dAb monotherapy and anti-CD28 dAb+MMF/steroids regimens are compared. Anti-CD28 dAb was compatible with traditional MMF/steroid immunosuppression and resulted in improved survival (MST >267 days) compared to monotherapy (MST >187 days). Addition of MMF/steroids was particularly beneficial in improving survival in the initial post-operative period with convergence of survival curves prior to 400 days post-transplant. One monkey in the dAb+ MMF/steroids treatment group maintained rejection-free survival at 600 days. The addition of T cell depletion to the anti-CD28dAb with conventional treatment regimen resulted no episodes of acute rejection during the therapeutic dosing period. (B) Drug levels of anti-CD28 dAb were consistent between NHPs receiving either anti-CD28 dAb monotherapy and those receiving anti-CD28 dAb with conventional MMF/steroids immunosuppression.

**Figure 4:**
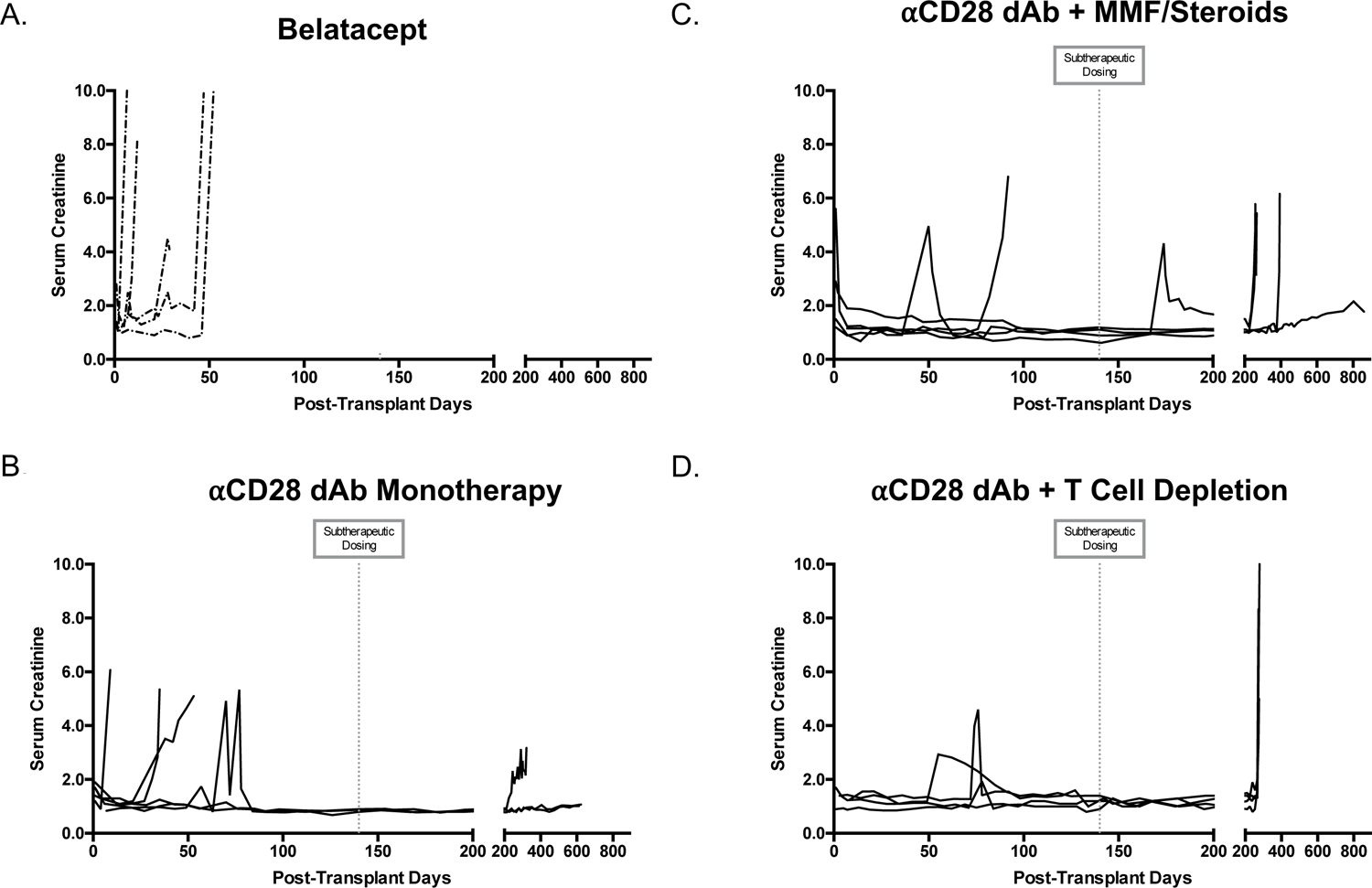
Serum creatinine levels for all treatment groups. (A) Belatacept control group. (B) Anti-CD28 dAb monotherapy (C) Anti-CD28 dAb+ MMF/steroids (D) Depletion-anti-CD28dAb+ conventional. Graft survival was associated with serum creatinine levels within a normal range.

**Table 1.** Treatment arms and survival times after NHP renal transplantation. Recipients were divided into four treatment arms: belatacept alone, anti-CD28 dAb alone, anti-CD28 dAb with anti-IL-2R mAb induction and anti-CD28 dAb with T cell depletion induction. Rejection-free survival times are listed as days post-transplant for all treatment groups. Rejection on therapy was defined as rejection occurring during the therapeutic portion of the dosing schedule. For anti-CD28 dAb this equaled rejection during weekly or biweekly injections (pre-POD 140). For belatacept this referred to rejection during biweekly or monthly infusions. Monkeys in the belatacept monotherapy group all rejected on therapy between POD 8-53. Survival times for animals treated with anti-CD28 dAb therapy alone ranged from 9 to >600 days. Three monkeys rejected during weekly injections of dAb. One monkey spaced to monthly treatment intervals at POD 140 and successfully remained on monthly injections for >180 days before rejecting at POD 321. Two animals maintained rejection-free survival >600 days; they were then sacrificed per protocol with no evidence of clinical or histologic rejection. Addition of conventional MMF/steroid therapy to anti-CD28 dAb resulted in survival from 92 to >600 days. Only one monkey in this treatment arm rejected on therapy. Two rejected once therapy was spaced from weekly/biweekly to monthly injections; however, they maintained rejection-free survival for >100 days on monthly treatment. The remaining monkey in this treatment group maintained graft tolerance with no rejection at >600 days. The depletion anti-CD28dAb with MMF/steroids treatment regimen was associated with no episodes of acute rejection during the therapeutic dosing period.

| Treatment Regimen | Survival (days) |
| --- | --- |
| **Belatacept**<br>MST 29 | 8*, 12*, 29*, 47*, 53* |
| **αCD28 dAb**<br>MST 187 | 9*, 35*, 53*, 321, >600, >600 |
| **αCD28 dAb + MMF/Steroids**<br>MST 267 | 92*, 259, 267, 394, >600 |
| **αCD28 dAb + T Cell depletion + MMF/Steroids**<br>MST 275 | 273, 274, 276, 280 |

\*Denotes rejection while on therapeutic dosing schedule

The addition of conventional maintenance immunosuppression (MMF/steroids) to anti-CD28 dAb improved early rejection rates and median survival from 187 to 267 days (*Table 1, Figure 3A*). In this treatment group, only one macaque rejected during biweekly anti-CD28 dAb injections. Two rejected after moving to subtherapeutic monthly injections, one rejected after cessation of drug therapy (discontinued at POD 364), and one showed no evidence of clinical or histologic rejection at 600 days (*Table 1*). The addition of conventional immunosuppression had no effect on serum drug levels compared to levels observed in animals treated with the anti-CD28 dAb alone (*Figure 3B*). Furthermore, survival in the anti-CD28 dAb plus conventional group appears to be superior than a similar belatacept plus conventional treatment regimen where median survival was 155 days (7). The addition of T cell depletion-based induction to the anti-CD28dAb and conventional treatment regimen was associated with no episodes of acute rejection during the therapeutic dosing period with (MST 275 days, *Table 1, Figure 3A*).

CMV titers were also assessed weekly post-transplant to ensure functional intact protective immunity against viral reactivation. No routine CMV prophylaxis was used, and antiviral therapy was initiated only for CMV viremia of >10,000 copies/mL. No cases of clinical CMV viremia were detected in anti-CD28 dAb or belatacept monotherapy groups. Addition of conventional therapy to anti-CD28 dAb resulted in one case of CMV viremia within the first 60 days post-transplant *(data not shown).* Viremia was easily treated with oral antiviral therapy with prompt resolution of peripheral viral load. Additionally, no clinical manifestations of CMV infection ensued. Overall, there does not appear to be a difference in the ability to maintain protective immunity between anti-CD28 dAb and belatacept.

### Treatment with anti-CD28 dAb increase circulating Tregs over time

In order to ensure that CD28-directed therapy did not impact T lymphocytic cellular subsets and to assess any changes in immune phenotype with CD28dAb treatment, peripheral blood was obtained weekly post-transplant for flow cytometric analysis. In recipients treated with anti-CD28 dAb monotherapy, CD4^+^ and CD8^+^ memory subsets maintained consistent frequencies at 30-weeks post-transplant (*Figure 5*). The improvements in graft survival with anti-CD28 dAb, particularly in the early post-transplant period, raise the question of whether intact inhibitory signaling through CTLA-4 contributes to this observation via a regulatory T cell dependent mechanism. To assess this, peripheral blood samples were examined for the frequency of CD25^+^ FoxP3^+^ CD4^+^ T cells. In order to determine if Treg frequencies correlated with the prolonged survival results in the anti-CD28 dAb-treated recipients, macaques were separated into “Rejectors” and “Non-rejectors” based on rejection status during the therapeutic-dosing treatment phase. Over time, recipients treated with anti-CD28dAb alone (N=3) or with conventional therapy (N=4) who did not reject during therapeutic drug dosing appear to experience a modest increase in frequency of Tregs in the periphery from pre-transplant values with a positive trend beginning around POD 90 and continuing up to 6-months post-transplant. (*Figure 6*). This data while not confirmatory or strictly causal does suggest that Treg numbers and possibly function was enhanced by anti-CD28 dAb treatment resulting in significant prolongation of kidney transplant survival even after completely weaning all immunosuppression.

**Figure 5.**
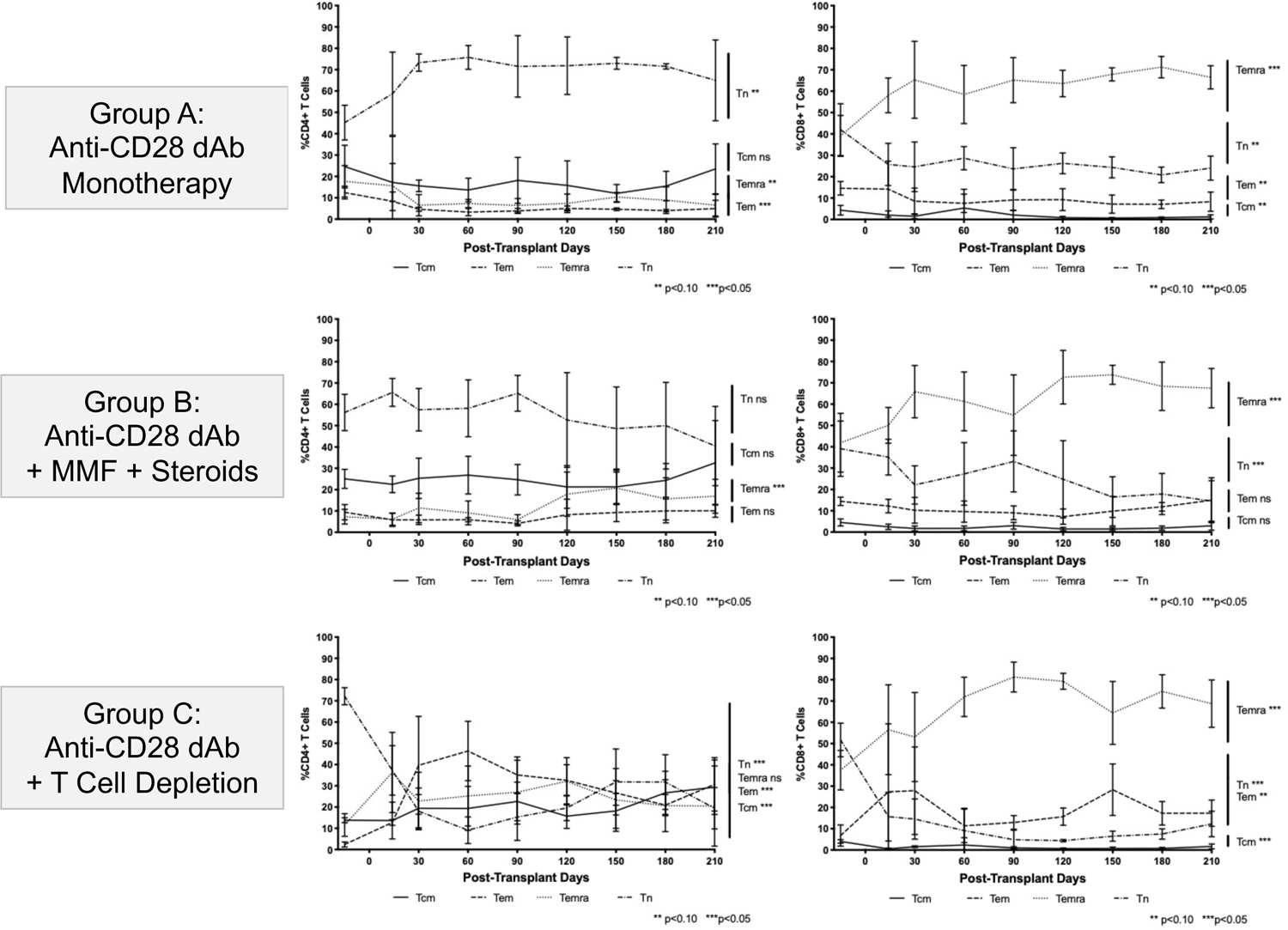
Regulatory T cells after use of costimulation blockade in renal transplantation. A. Peripheral blood immunophenotypic analysis was performed using flow cytometry. After gating on CD3^+^CD4^+^ T cells, CD25^+^Foxp3^+^ cells were selected as Tregs. Beginning at POD 90, peripheral Treg counts in non-rejectors were significantly increased. (B) Representative flow cytometric analysis of Treg frequencies for a recipient treated with anti-CD28 dAb-treated POD154 is shown.

**Figure 6.**
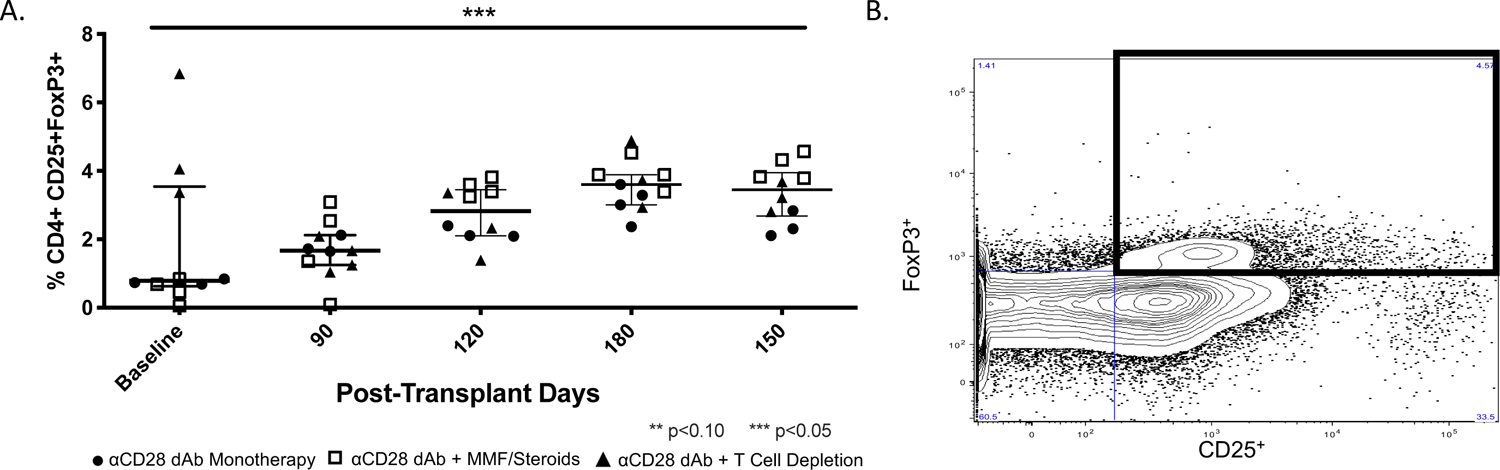
Development of DSA after treatment with anti-CD28 dAb. Donor specific antibody (DSA) production was evaluated serially post-transplant and binding of antibody to both Class I and Class II MHC was assessed. Time of DSA development refers to the POD when an animal tested positive for DSA. In each case, the animal was assessed at previous time points (generally every 1-2 months) and was negative. No recipients developed post-transplant DSA on therapeutic dosing. Two monkeys developed DSA once treatment intervals increased to monthly injections, and three additional recipients developed DSA once off therapy. DSA in two of these recipients was directed against Class II MHC. No clinical or histologic evidence of rejection was seen in these recipients despite the presence of DSA in serum.

### Therapeutic doses of anti-CD28 dAb prevent the development of donor-specific antibodies

We did not observe the development of donor specific antibody in the serum of animals when sampled during therapeutic anti-CD28 dAb dosing, i.e., weekly or biweekly injections. After treatment was spaced to monthly intervals, two animals developed donor-specific antibodies. Three additional animals exhibited anti-donor antibody levels after therapy was halted (*Figure 7*). Antibodies were primarily directed against MHC Class II. These data further suggest that weekly or biweekly subcutaneous dosing is sufficient to prevent renal graft rejection including antibody-mediated rejection. Belatacept, which is known to have lower rates of donor specific antibody formation than standard immunosuppression with calcineurin-based immunosuppression, did not allow the development of donor specific antibodies in animals on therapy either.

**Figure 7.**
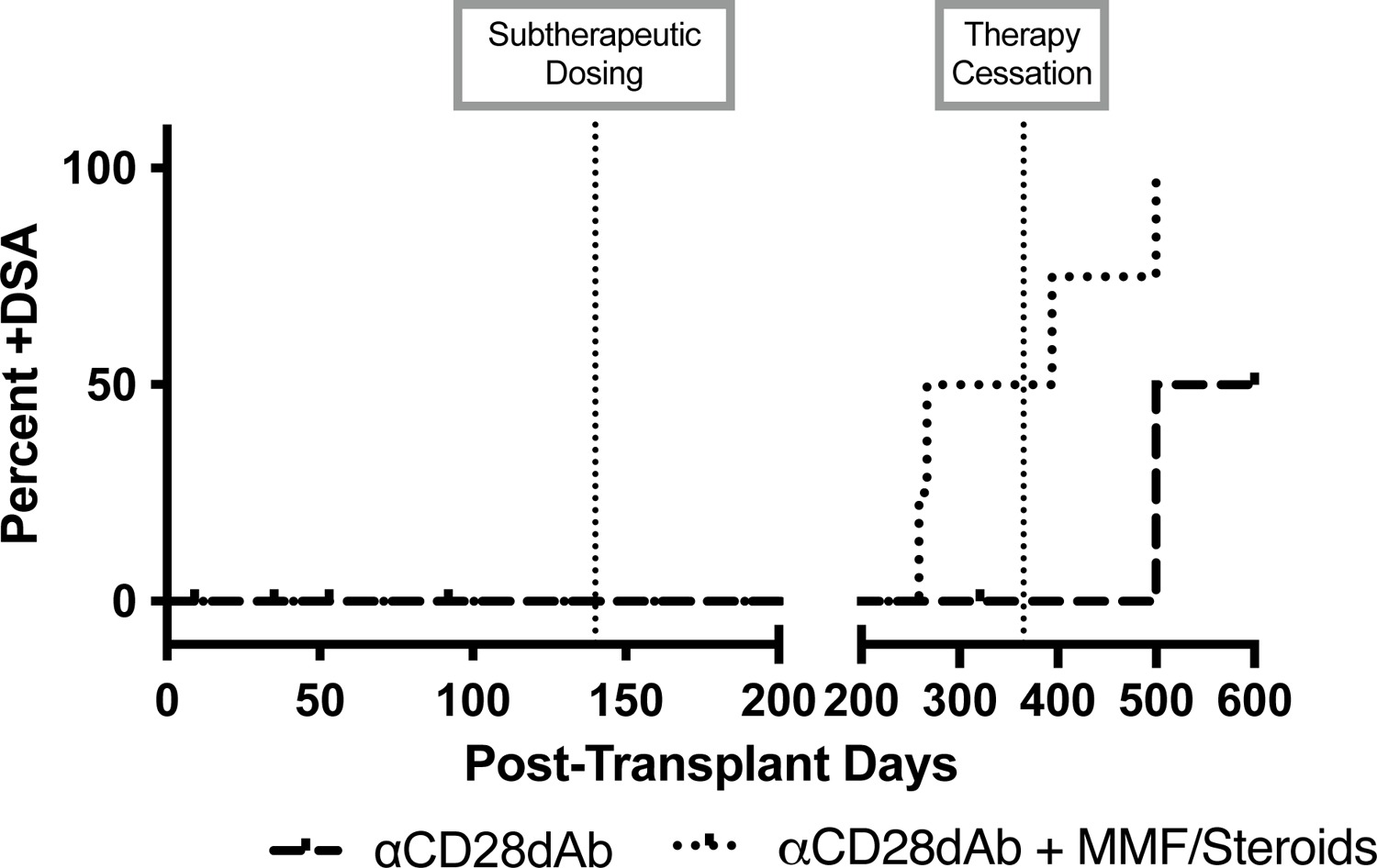

## Discussion

Renal transplantation improves survival and quality of life for thousands of patients suffering from renal failure. The development of novel targeted immunosuppressive therapies offers the ability to extend this benefit to further improve allograft and patient survival. Standard immunosuppressive regimens containing CNI have significant limitations and while belatacept has been shown in a randomized, clinical trial to improve survival and renal function compared to cyclosporine, the first post-transplant year is marked by increased rates of acute rejection necessitating many patients continue on low-dose CNI for some period of months before weaning (20). Furthermore, belatacept requires intravenous administration often resulting in significant inconvenience to patients. A superior approach would be to target CD28 directly, preventing that key costimulatory signal while preserving co-inhibitory signaling through CTLA-4 unlike belatacept which blocks both signals. Additionally, anti-CD28 dAb is injected subcutaneously and gives patients the flexibility of self-administration providing a more attractive and practical approach. Furthermore, anti-CD28 dAb was well tolerated and demonstrated a favorable safety profile in a phase II clinical trial of patients with systemic lupus erythematosus (SLE) (NCT02265744). The current study demonstrated that a CD28-selective approach to immunosuppression can prevent rejection in a non-human primate model of renal transplantation.

Anti-CD28 dAb improves long-term survival with low rates of early rejection in our preclinical model of renal allograft transplantation in rhesus macaques. When anti-CD28 dAb monotherapy was directly compared to belatacept the data suggested an improvement in long-term survival with anti-CD28 dAb although it did not achieve statistical significance. While median survival rates were similar between monotherapy groups, we did observe the development of long-term therapy-free survival in recipients treated with anti-CD28 dAb but not in animals treated with belatacept. This finding of long-term therapy free survival was also observed in an anti-CD28 dAb treatment group who received additional conventional therapy.

While every monkey receiving monotherapy belatacept rejected prior to POD 60 while on therapy, only half of the anti-CD28 dAb monkeys rejected during this same time frame. For monkeys who survived the early post-transplant phase, rejection did not occur until the dosing interval was increased to monthly injections, where drug reached subtherapeutic levels. The addition of conventional immunosuppressive agents (MMF and steroids) to anti-CD28 dAb helped to eliminate early post-transplant rejection. Only one out of six recipients in the anti-CD28 dAb with conventional immunosuppression rejected while on biweekly therapy, compared to three out of eight belatacept treated animals in a control group. Importantly, all anti-CD28 dAb-containing regimens in this study were free of any CNI. Previous groups testing CD28-directed therapies have required low-doses of CNI to obtain survival greater than median 18.5 days (15), suggesting a reliance on CNI to control the alloimmune response. Data in this study also suggests that weekly or biweekly subcutaneous injections of anti-CD28 dAb, in combination with conventional immunosuppression, is sufficient to avoid early post-transplant rejection, and may give rise to long-term, therapy free survival. Furthermore, no donor specific antibody developed until recipients reached subtherapeutic dosing with biweekly treatment intervals for at least 140 days. The addition of pretransplant T cell depletion was associated with a relative freedom from rejection in the early posttransplant period, including no episodes of rejection during therapeutic CD28dAb treatment suggesting the T cell depletion-based induction might be the best strategy to avoid early rejection when using anti-CD28dAb.

The improvements in graft survival achieved with anti-CD28 dAb did not occur with a tradeoff for safety or loss of protective immunity. While a previous iteration of CD28-selective therapy, TGN1412, resulted in dramatic lymphopenia (18), no CD4^+^ or CD8^+^ T cell depletion occurred with the anti-CD28 dAb in this study. Furthermore, all memory T lymphocyte subsets remained stable post-transplant, suggesting a favorable safety profile in NHP. No cases of CMV reactivation occurred in the monotherapy regimen, and only one animal had CMV reactivation in the anti-CD28 dAb plus conventional immunosuppression group, similar to the belatacept treated control group suggesting that protective immunity is maintained during anti-CD28 dAb treatment.

To find a mechanistic explanation for improved survival, differences in peripheral blood T cell subsets were assessed between anti-CD28 dAb and belatacept. Treg frequencies were assessed for higher numbers in the anti-CD28 dAb-treated recipients, given the known importance of CTLA-4 for proper Treg functioning and since previous work with the CD28-directed therapy FR104 suggested possible increases in Tregs post-transplant (15). We found that treatment with anti-CD28 dAb also may lead to increased Tregs in Rhesus macaques over time. However, the increase was not notable until 90 days post-transplant. Given the relatively late increase in Tregs, this may not explain the differences in early rejection rates between anti-CD28 dAb and belatacept. Additionally, the trend of increasing Tregs noted at later timepoints may be due to positive selection among long-term survivors.

Benefits of this pre-clinical renal transplant investigation include use of a biologic agent already shown in phase II clinical trial to have a favorable safety profile. An additional benefit includes the ability to test anti-CD28 dAb in a rigorous NHP model with maximally MHC-mismatched donor/recipient pairings. Limitations of the current study include the relatively small number of recipients in each treatment group. This may be contributed to difficulty in obtaining significant results in the survival data; however, the trend towards improved survival with anti-CD28 dAb remains clear. In conclusion, CD28-directed therapy alone or in combination with conventional immunosuppression prolongs renal allograft survival in a preclinical renal allograft model with non-human primates and is superior to belatacept. Anti-CD28 dAb therapy represents a superior alternative to belatacept as a strategy to promote CNI-free immunosuppression that presumably would promote improved long-term survival with increased renal function and less rejection.

## Acknowledgements

This work was supported by the Non-Human Primate Transplant Tolerance Cooperative Study Group, NIAID U19AI051731 and NIH grant P51OD11132. L.H. and S.C.K. are supported by an NIH Ruth L. Kirschstein NRSA Institutional Research Training Grant (T32). The anti-CD28 dAB was provided by Bristol Myers Squibb.

---

**Supplemental Figure 1.**
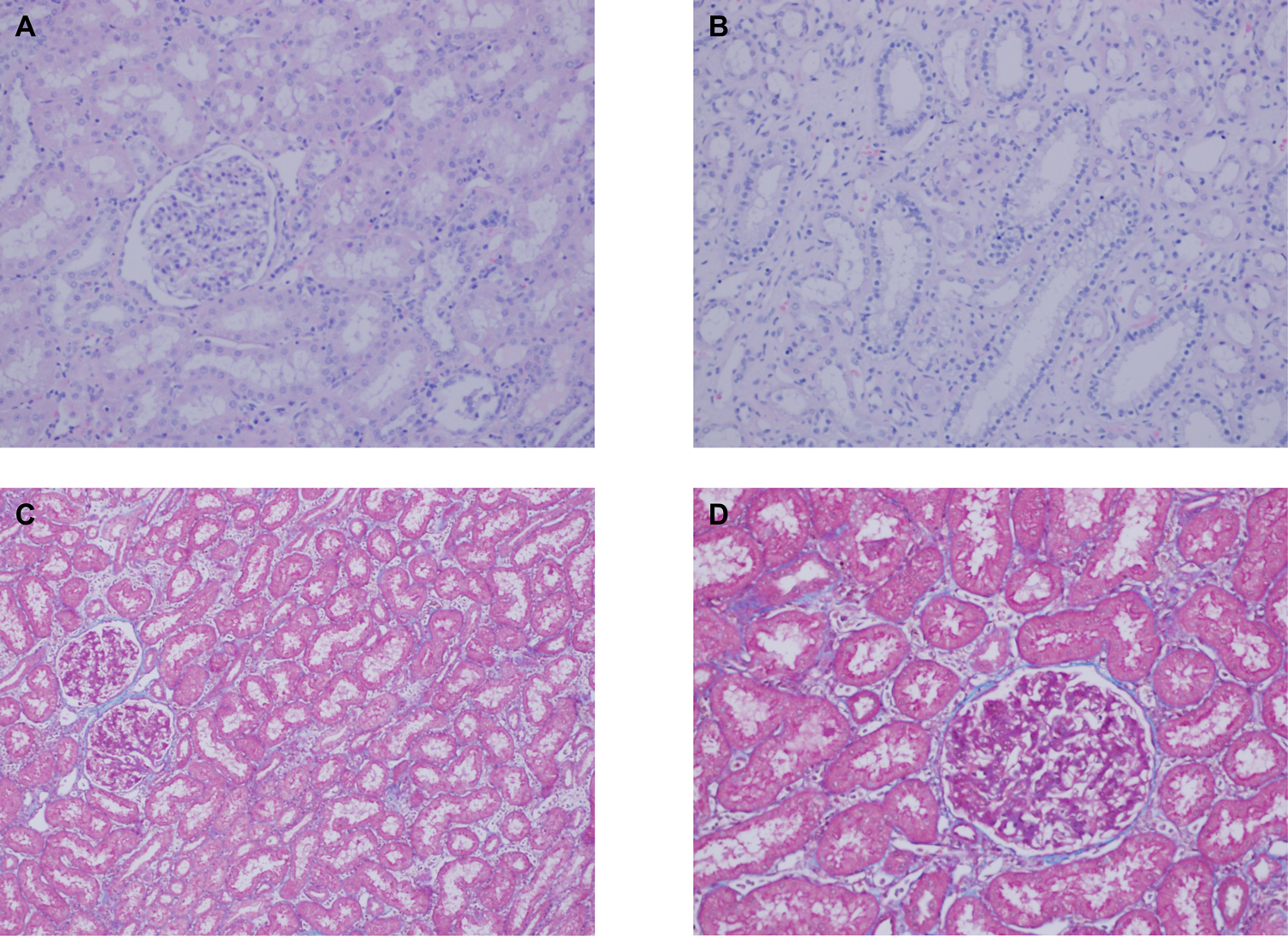
Histologic analysis. Monkeys who remained ongoing at 600 days with no clinical evidence of rejection (n=3) were sacrificed for retrieval of the transplanted renal graft. Specimens were submitted for H&E and trichrome staining with representative images shown. H&E showed minimal evidence of cellular infiltrate in (A) glomeruli and (B) tubules. (C-D) Trichrome demonstrated no significant interstitial fibrosis and no tubular atrophy.

